# Carbon Fiber Electrodes for Intracellular Recording and Stimulation

**DOI:** 10.1101/2021.08.12.456117

**Authors:** Yu Huan, Jeffrey P. Gill, Johanna B. Fritzinger, Paras R. Patel, Julianna M. Richie, Elena della Valle, James D. Weiland, Cynthia A. Chestek, Hillel J. Chiel

## Abstract

To understand neural circuit dynamics, it is critical to manipulate and record from many neurons, ideally at the single neuron level. Traditional recording methods, such as glass microelectrodes, can only control a small number of neurons. More recently, devices with high electrode density have been developed, but few of them can be used for intracellular recording or stimulation in intact nervous systems, rather than on neuronal cultures. Carbon fiber electrodes (CFEs) are 8 micron-diameter electrodes that can be organized into arrays with pitches as low as 80 µm. They have been shown to have good signal-to-noise ratios (SNRs) and are capable of stable extracellular recording during both acute and chronic implantation *in vivo* in neural tissue such as rat motor cortex. Given the small fiber size, it is possible that they could be used in arrays for intracellular stimulation. We tested this using the large identified and electrically compact neurons of the marine mollusk *Aplysia californica*. The cell bodies of neurons in *Aplysia* range in size from 30 to over 250 µm. We compared the efficacy of CFEs to glass microelectrodes by impaling the same neuron’s cell body with both electrodes and connecting them to a DC coupled amplifier. We observed that intracellular waveforms were essentially identical, but the amplitude and SNR in the CFE were lower than in the glass microelectrode. CFE arrays could record from 3 to 8 neurons simultaneously for many hours, and many of these recordings were intracellular as shown by recording from the same neuron using a glass microelectrode. Stimulating through CFEs coated with platinum-iridium had stable impedances over many hours. CFEs not within neurons could record local extracellular activity. Despite the lower SNR, the CFEs could record synaptic potentials. Thus, the stability for multi-channel recording and the ability to stimulate and record intracellularly make CFEs a powerful new technology for studying neural circuit dynamics.

## 1. Introduction

Clarifying the dynamics of neural circuitry continues to be a major challenge for neuroscience, and developing new technologies for monitoring and manipulating neural activity will be critical for advances in the field. Ideally, a technique for studying a neural circuit should have several features. First, the technique needs to be able to record simultaneously from large numbers of neurons. Second, the technique should monitor intracellular potentials, including the subthreshold membrane potentials of individual neurons, so that synaptic connections and their role in controlling neural activity can be clarified. Third, it should be possible to implant the recording and stimulating device in intact, behaving animals, and generate stable long-term recordings. Finally, the device should both be able to record and inhibit or excite neurons to determine the causal role of individual neurons or groups of neurons in circuit function.

Obtaining stable long-term intracellular interfaces for recording and stimulation is particularly challenging. In general, intracellular electrodes penetrate the cell membrane, which could cause damage to the neuron, especially if an animal moves. The recording devices also need to have appropriate spacing to monitor as many adjacent neurons as possible without multiple penetrations of a single cell. Despite these difficulties, intracellular recordings are critical because subthreshold synaptic activity serves important physiological functions in a neural network [1-2].

Current intracellular techniques do not meet these requirements. The sharp glass microelectrode has been a traditional tool for many years [3], and provides the ability to completely control a neuron’s membrane potential and to monitor sub-threshold activity. However, an electrode is restricted to a single neuron, and will damage the neuron if the preparation moves. Voltage- and calcium-sensitive dyes can record from many neurons simultaneously and are non-invasive [4], but may induce pharmacological effects and require computational methods to faciliate signal intepretation [5-6]. Genetically-encoded voltage sensors show great promise [7-8], but they require genetic manipulation and high quality imaging equipment to achieve high resolution, and cannot yet be used in freely-behaving animals. Other novel electrode-like intracellular techniques have been developed, but they either have not been applied to a large number of neurons for stimulation or have limited recording stability [9-11]. Furthermore, these electrode-like intracellular techniques may be difficult to use in intact, freely-behaving animals. Improvements in devices will be needed to investigate neural network dynamics during natural, unconstrained behavior.

Carbon fiber electrodes (CFEs) are a relatively new technique that has been developed and improved over time [12-14]. Earlier work used carbon paste fibers for voltammetry [15], but the fibers are large (50 µm – 1.6 mm diameter) and thus not suitable for recording from individual neurons. In more recent work on CFEs, individual carbon fibers have diameters of about 8.4 µm (including Parylene C insulation), a good signal-to-noise ratio (SNR), and can generate stable extracellular recordings chronically *in vivo* [16]. The fibers can be arranged in arrays of 16 electrodes with interelectrode spacing from 80 to 150 µm. The small diameter of the electrodes suggests that it might be possible to use them for intracellular stimulation and recording. Previous work in songbird auditory forebrain nuclei reported intracellular-like action potentials recorded by small diameter carbon fiber electrodes (∼ 5.5 µm including Parylene C insulation) [12], but this was not fully explored. Tests for intracellular recording and stimulation are still needed.

A suitable test subject for determining whether CFEs can be used for intracellular recording and stimulation should have large neurons that are electrically compact, and have well-defined synaptic interactions. This has been our rationale for testing the intracellular use of CFEs in the marine mollusk *Aplysia californica* [17-18]. Motor neurons are often about 100 µm in diameter, and thus well-matched to the pitch of CFEs. In addition, *Aplysia*’s neurons are electrically compact, and many details of the synaptic interactions between neurons have been intensively studied, making them ideal for testing the ability of CFEs to intracellularly record and stimulate. In particular, in the collection of nerve cells that control feeding behavior in *Aplysia*, the buccal ganglion, the relationship between a multi-action neuron (B4/B5) and its synaptic followers has been very well-characterized [19].

In this study, we investigated the effectiveness and stability of CFEs for intracellular recording and stimulation. By inserting a CFE and a traditional glass microelectrode into the same neuron to directly compare the two kinds of electrodes, we found that the CFE could measure subthreshold membrane potentials and action potentials almost as well as a glass microelectrode. We also measured the recording yield and the current needed for stimulation using CFEs. Current injected through CFEs successfully excited or inhibited neurons, suggesting that this new device could be a new and effective approach to monitoring and manipulating neural circuitry.

## 2. Materials and Methods

### 2.1 Animals

*Aplysia californica* were acquired from South Coast Bio-Marine (San Pedro, CA) or Marinus Scientific (Newport Beach, CA) and kept in aerated aquaria containing artificial seawater at 15.5°C on a 12/12 hour light/dark cycle. Animals of 100 - 350 g were used.

Animals were anethesized using an injection of 333 mM isotonic magnesium chloride solution in a volume half of the animals’ body weight [20]. The buccal mass was dissected out and hook electrodes were attached to buccal nerves (details below, section 2.3). The buccal ganglia were then cut away from the buccal mass. The isolated buccal ganglia were placed in a Petri dish and pinned to a Sylgard base using insect pins. The sheath of the buccal ganglion ipsilateral to the recording hook electrodes was completely removed to expose the neurons (Figure 1 B, D, E) in a solution that was half *Aplysia* saline (460 mM NaCl, 10 mM KCl, 22 mM MgCl_2_, 33 mM MgSO_4_, 10 mM CaCl_2_, 10 mM glucose, and 10 mM MOPS, pH 7.5) and half isotonic magnesium chloride to minimize movement of the sheath during dissection. During the recordings, to maintain normal neural activity, the ganglia were kept in normal *Aplysia* saline; to inhibit polysynaptic transmission between neurons, ganglia were exposed to a high divalent cation solution (270 mM NaCl, 6 mM KCl, 120 mM MgCl_2_, 33 mM MgSO_4_, 30 mM CaCl_2_, 10 mM glucose, and 10 mM MOPS, pH 7.5); finally, to evoke spontaneous neural activity, ganglia were exposed to a high potassium solution (420 mM NaCl, 50 mM KCl, 22 mM MgCl_2_, 33 mM MgSO_4_, 10 mM CaCl_2_, 10 mM glucose, and 5 mM MOPS, pH 7.5).

**Figure 1.**
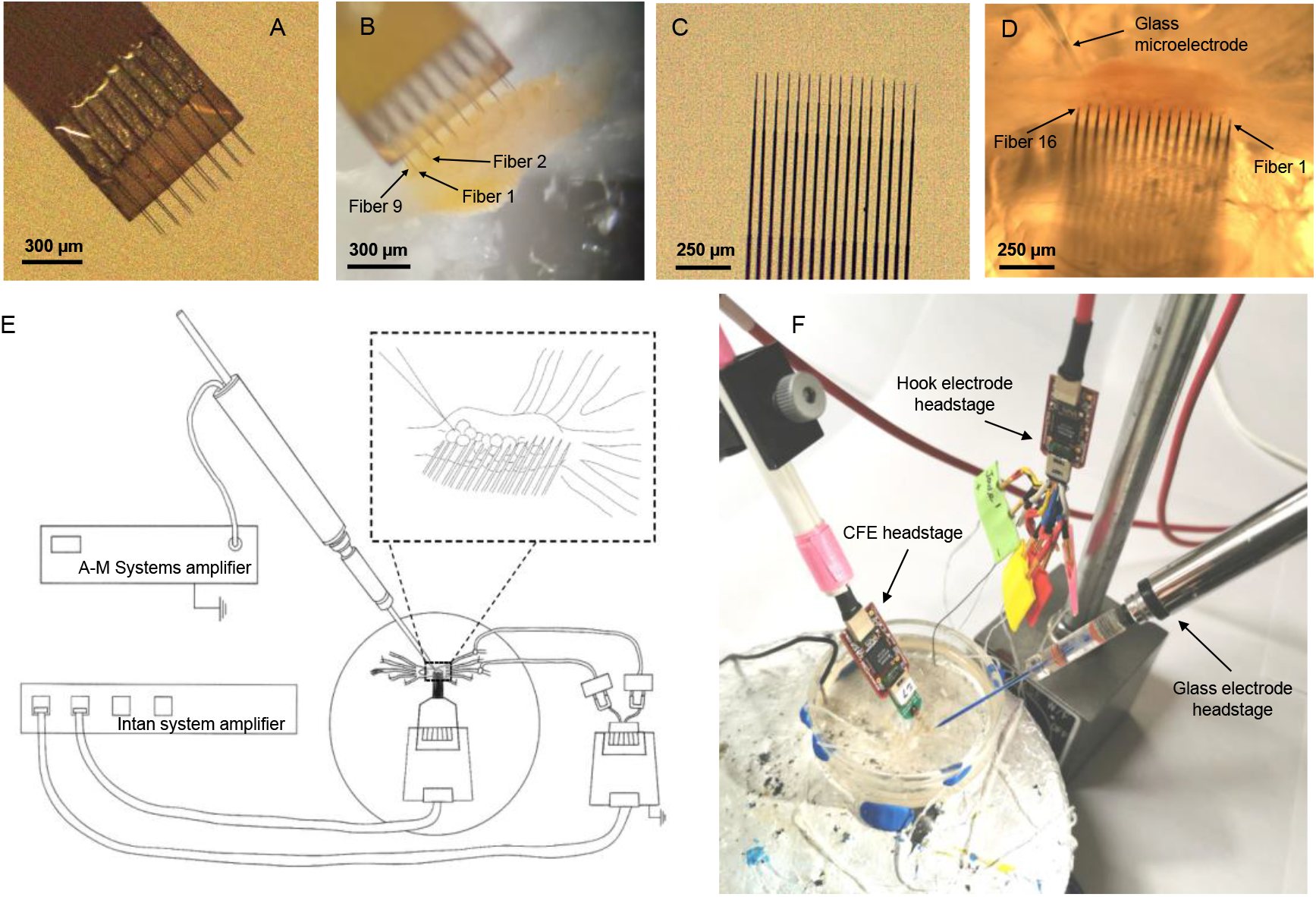
Carbon fiber electrode arrays for recording and stimulation. (A) A flex array. The fibers were sharpened and coated with PEDOT:pTS at the tip. See text. (B) A representative photograph of a flex array inserted into a buccal ganglion. (C) A high density carbon fiber (HDCF) array. The fibers were sharpened and coated with either PEDOT:pTS or platinum-iridium (PtIr) at the tip. (D) A representative photograph of an HDCF array inserted into a buccal ganglion along with a glass microelectrode. (E) A schematic of the experimental setup. The nerve recording headstage and the CFE recording headstage were connected to the Intan amplifier system (see text). The glass microelectrode was connected to a DC-coupled AM Systems amplifier for intracellular recordings. Inset shows a closer view of the relationship between the carbon fibers and the neurons in the ganglion. (F) A photograph of the experimental setup. The glass microelectrode and the CFE were positioned close to the buccal ganglion, and different headstages were connected to the CFEs and to the glass microelectrode.

### 2.2 Carbon Fiber Electrode Fabrication

Carbon fiber electrode (CFE) arrays were fabricated in the laboratory of Dr. Chestek at the University of Michigan. Two arrays with different configuration were used: a flex array (Figure 1 A, B) and a high density carbon fiber (HDCF) array (Figure 1 C, D). The flex array has a two by eight configuration, with a 132 µm pitch. Detailed fabrication instructions for the flex array can be found in [21-22]. The HDCF array (Figure 2 A) has a one by sixteen configuration, with a 100 µm pitch. This array consists of a minimally invasive silicon support structure (see below) that provides a permanent shuttle for the carbon fiber electrodes.

**Figure 2.**
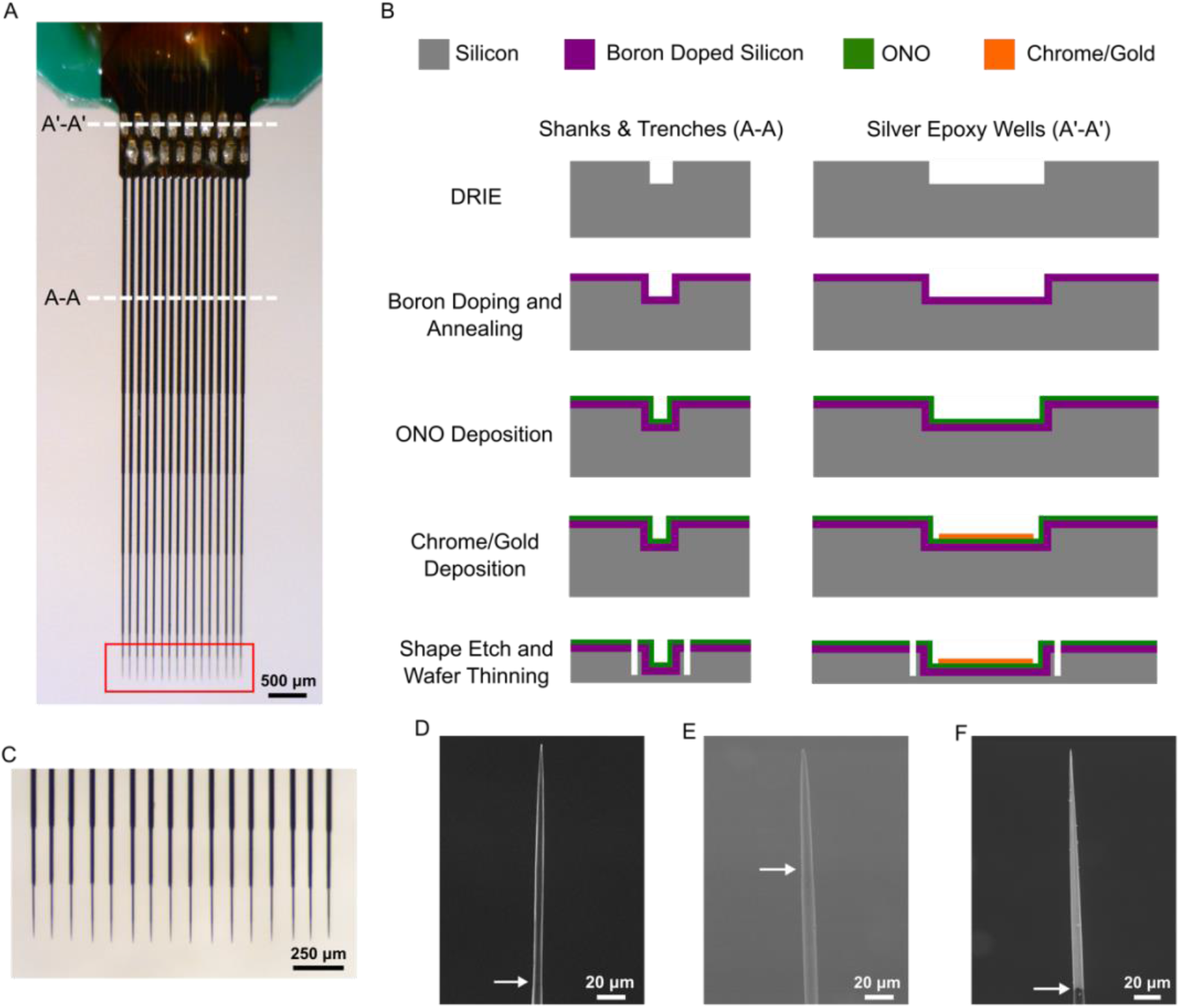
HDCF array fabrication. (A) Image of a fully assembled and populated HDCF array. Red box indicates region of array that was blowtorch sharpened (see text). (B) Illustration of the cleanroom fabrication steps for the silicon support structure. Briefly, a deep reaction ion etching (DRIE) step creates the trenches for the fibers (A-A) and silver epoxy wells (A′-A′). Boron doping and annealing creates an eventual etch stop for the last release step. Deposition of ONO (see text) creates an insulation layer, on top of which chrome and gold are deposited and patterned in the silver epoxy wells. Lastly, the overall device shape is achieved with another DRIE step, the backside thinned, and final release is achieved with an EDP wet etch. (C) Close up from (A) showing approximately 250 µm of carbon fiber protruding from the ends of the silicon supports. The very tips of the CFEs have been blowtorch sharpened. (D) SEM image of a blowtorch sharpened CFE. (E) SEM image of a sharpened CFE with a PEDOT:pTS coating. (F) SEM image of a sharpened CFE with a PtIr coating. Arrows for all SEM images indicate the transition from Parylene C to the bare or coated portion of the CFE.

#### 2.2.1 High Density Carbon Fiber Array - Silicon Support Structure Fabrication

The fabrication of the silicon support structure (Figure 2 B) started by deep reaction ion etching (DRIE) (STS Pegasus 4; SPTS Technologies, Newport, United Kingdom) of a 4” silicon wafer (P-10-20; Silicon Valley Microelectronics, Inc., Santa Clara, CA) to form both the silver epoxy wells and shank trenches where the fibers would be placed. The overall length of the trenches defined the length of the device, which in this application is 6 mm. Next, the wafers were boron doped and annealed, which provides for an etch stop during the final release step. After annealing, the wafer underwent a low pressure chemical vapor deposition of high temperature oxide – nitride – high temperature oxide (ONO). Layer thicknesses, 1500 Å – 463 Å - 1500 Å, were chosen to cancel the compressive and tensile stresses introduced by the oxide and nitride films, respectively.

Next, a chrome adhesion layer (t = 300 Å) and gold layer (t = 3000 Å) were sputter deposited (Lab 18 Sputtering System, Kurt J. Lesker, Jefferson Hills, PA) and then wet etched onto the silver epoxy wells. These layers created an electrical contact between the silver epoxy wells and pads that would eventually be wire bonded to an external printed circuit board. The support structured the final shape, including the tapering of the shanks, which was defined by another DRIE step. Before the final wet etch release, the backside of the wafers underwent a series of DRIE steps to remove approximately 450 to 500 µm of silicon. To remove the remaining un-doped silicon and release the device an ethylenediamine pyrocatechol (EDP) wet etch was used, which has high selectivity against boron doped silicon.

#### 2.2.2 High Density Carbon Fiber Array – Assembly

To begin, a connector (A79040-001; Omnetics, Minneapolis, MN) was soldered to a custom printed circuit board (PCB). The pins of the connector were then covered with two-part epoxy (Sy-SS; Super Glue Corporation, Ontario, CA). Next, the silicon support structure was secured to the PCB using epoxy (301; Epoxy Technology, Billerica, MA) cured at 140 °C for 20 minutes. The overhanging, underside portion of the silicon support that consisted of the silver epoxy wells was reinforced to the PCB with two-part epoxy (Sy-SS; Super Glue Corporation, Ontario, CA). Devices were then wire bonded to connect the gold pads on the silicon supports to the pads on the PCB. The wire bonds were then covered with another epoxy (353NDT; Epoxy Technology, Billerica, MA) cured at 140 °C for 20 minutes.

To place the individual carbon fibers, first a droplet of epoxy (NOA 61; Norland Products, Inc., Cranbury, NJ) was briefly held at the tips of the support to allow for a small amount to wick up each shank approximately one-third of the way. At the other end (silver epoxy wells), a small amount of deionized water was deposited, which also flowed along the trenches and stopped at the epoxy. Then, individual carbon fibers (T-650/35 3K; Cytec Thornel, Woodland Park, NJ) were cut to length (∼ 9 mm) and manually placed in the trenches using forceps. Care was taken to ensure that at least half the length of the silver epoxy well was occupied by each fiber. The epoxy was cured in an UV oven (ZETA 7401; Loctite, Westlake, OH) for 2 minutes.

Silver epoxy (H20E; Epoxy Technology, Billerica, MA) was deposited in the silver epoxy wells using an NLP 2000 system (Advanced Creative Solutions Technology, Carlsbad, CA). The epoxy was cured at 140 °C for 20 minutes. The exposed gold-silver epoxy-carbon fiber bond was then covered with epoxy (NOA 61; Norland Products, Inc., Cranbury, NJ) and additional NOA 61 epoxy was applied along the shanks to fully secure the carbon fibers before UV curing. Fibers were then cut to approximately 300 to 350 µm and coated with approximately 800 nm of Parylene C (PDS 2035; Specialty Coating Systems, Indianapolis, IN), before final tip functionalization.

#### 2.2.3 Tip Functionalization and Reference/Ground Wires

To functionalize the carbon fiber tips, regardless of array type, the CFEs were first blowtorch sharpened (Figure 2 A (red box), C, D) following methods described in [23]. Next, one of two materials (PEDOT:pTS or Platinum-iridium) was electrodeposited.

The first started with a mixture of 0.01 M 3,4-ethylenedioxythiophene (483028; Sigma-Aldrich, St. Louis, MO):0.1 M sodium p-toluenesulfonate (152536; Sigma-Aldrich, St. Louis, MO). Electrodeposition of this solution was carried out by applying 600 pA/fiber for 600 s to form a layer of poly(3,4-ethylene dioxythiophene):sodium p-toluenesulfonate (PEDOT:pTS) (Figure 2 E) [16, 23].

Prior to platinum-iridium (PtIr) plating, the CFEs underwent plasma ashing using a Glen 1000P Plasma Cleaner (pressure 200 mT, power 300 W, time 120 s, oxygen flow rate 60 sccm, and argon flow rate 7 sccm). To plate, a solution of 0.2 g/L of Na_3_IrCl_6_H_2_O (288160; Sigma-Aldrich, St. Louis, MO) and 0.186 g/L of Na_2_PtCl_6_H_2_O (288152; Sigma-Aldrich, St. Louis, MO) in 0.1 M of nitric acid (438073; Sigma-Aldrich, St. Louis, MO) was used (Figure 2 F) [24]. The solution was boiled until the color became reddish and was then cooled down to room temperature. A 70 μm PtIr wire (778000; A-M Systems, Sequim, WA) electrode was used as a counter electrode and an Ag|AgCl electrode as the reference (RE-5B; BASi, West Lafayette, IN). The potential range was set between -0.1 to 0.1 V with a scan rate of 200 mV/s for 1200 cycles, which corresponds to a coating process time of 45 minutes. The coating temperature was set to 56 °C and pulsed sonication at a power of 2 W (T_ON_ = 1 min and T_OFF_ = 30 sec.) was used to improve the coating rate. A Gamry 600+ potentiostat (Gamry Instruments, Warminster, PA) was used to apply potential cycles and an A700 Qsonica (Qsonica L.L.C., Newtown, CT) sonicator was used for sonication.

After tip functionalization, silver reference and ground wires (AGT05100; World Precision Instrument, Sarasota, FL) were attached to the PCB.

#### 2.2.4 Scanning Electron Microscopy Imaging

Scanning electron microscopy (SEM) images of the carbon fiber electrodes were acquired using a Tescan Rise SEM (Tescan Orsay Holding, Brno - Kohoutovice, Czech Republic) in low vacuum mode with an excitation voltage between 5 - 10 kV. The low vacuum mode allows for imaging without the deposition of a conductive film (e.g. gold).

### 2.3 Glass and Hook Electrode Fabrication

Intracellular glass microelectrodes were prepared to directly compare results from the CFEs. They were made from glass capillary tubes with a filament (615000; A-M Systems, Everett, WA) pulled by a Flaming–Brown micropipette puller (P-80/PC; Sutter Instruments, Novato, CA) [25-26]. Intracellular electrodes were backfilled with 3 M potassium acetate. To confirm an intracellular recording by the CFE for multiple neurons and to visualize the insertion site, several crystals of the dye Fast Green FCF (F7258; Sigma, St. Louis, MO) were added to the potassium acetate as the electrode was backfilled. The impedances of the intracellular micropipettes ranged between 2.5 - 6 MΩ.

Extracellular hook electrodes were prepared to record nerve activity during motor patterns, and to confirm that when neurons were activated by the CFEs at the soma, this activation induced propagating action potentials in the axons of the neurons that then propagated through the nerves. They were prepared as described by [20] (see section 3, steps 3-13). Briefly, hook electrodes were made from enamel-insulated stainless steel 316 wire (100194, 25 μm diameter, heavy polyimide insulated; California Fine Wire Company, Grover Beach, CA). Two wires were coated with silicone glue to make a single-channel twisted pair. The silicone glue and the enamel on both ends of both wires were stripped away to expose the electrically conductive wire. At one end, each wire was soldered to a male gold pin. On the other end, one wire was bent into a hook-like shape to be placed around a nerve whereas the other wire was used as a reference wire.

During the experiments, the hook electrodes were attached to buccal nerve 2 (BN2) and buccal nerve 3 (BN3) ipsilaterally for recording, because most of the key motor neurons project through these nerves [27]. A stimulating hook electrode was also attached to contralateral buccal nerve 2-a (BN2-a; [28], a sensory branch of buccal nerve 2, to trigger motor patterns, which helps to identify neurons [27].

### 2.4 CFE Experiments using an Intracellular Amplifier

The CFEs were first tested using an intracellular amplifier (Neuroprobe Amplifier Model 1600; A-M Systems, Everett, WA). The A-M Systems amplifier is DC-coupled and provides an accurate measurement of intracellular membrane potentials. However, since the intracellular A-M Systems amplifier is designed for single channel recordings, a single CFE from a flex array was used for the test.

During the experiments, a glass microelectrode and a CFE were inserted into the soma of the same neuron and each was connected to its own intracellular A-M Systems amplifier. The hook electrodes that recorded nerve signals were connected to an extracellular amplifier (Differential AC Amplifier Model 1700; A-M Systems, Everett, WA) to monitor nerve activity. Recordings were obtained simultaneously in AxoGraph X (AxoGraph Scientific, Foster City, CA) at a sampling frequency of 10 kHz.

The sharp glass microelectrode was held by an electrode holder (671440; A-M Systems, Everett, WA), which transmitted signals to the A-M Systems amplifier through a headstage (681500; A-M Systems, Everett, WA). A CFE was connected to the same type of headstage through a customized connector. This connector was made by soldering a male pin connector (521000; A-M Systems, Everett, WA) to a female nano-strip connector (A79025-001; Omnetics, Minneapolis, MN) so that it could interface with the A-M Systems intracellular amplifier. Silicone glue (GE284, ASTM C920 Class 35; GE Silicone, Rocky Hill, CT) was applied around the male pin connector to reduce signal drift and noise in the recording. Both headstages were held by hydraulic micromanipulators (MO-203; Narishige, Tokyo, Japan), which allowed fine control of movement of both electrodes.

After desheathing the buccal ganglion and setting up the two headstages, the CFE was first inserted into the neuron’s soma, followed by the insertion of the glass microelectrode into the same soma. Since the CFE has a certain flexibility, inserting it first minimized the damage that would be done to the cell membrane by the hard tip of a glass microelectrode. After determining the suprathreshold stimulating current for the CFE, identical monophasic excitatory currents were injected alternately into each electrode to evoke action potentials which could be recorded by the other electrode.

In other experiments, minimum currents that could inhibit spontaneous activity (in high potassium saline or in normal *Aplysia* saline) or excite the neuron (in normal *Aplysia* saline) were first determined for the CFE, and then the same current was injected through the glass microelectrode.

### 2.5 CFE Experiments using Extracellular Amplifiers

To obtain multiple simultaneous recordings from the entire array of CFEs and to stimulate from multiple CFEs, they were connected to an Intan RHS 32-channel system (M4200; Intan Technologies, Los Angeles, CA). The Intan system is AC-coupled with built-in analog filters and so cannot provide DC-coupled recordings, but the filters can be set so that near-DC recordings can be obtained. The array of CFEs was designed to be compatible with the Intan system headstage, so no customized connector was required.

The CFEs and the hook electrodes were both connected to the Intan system and recorded using the Intan stimulation/recording controller software. The cutoff frequencies of the two AC amplifiers could be adjusted through the software. During the data acquisition, the low-pass cutoff frequency was set to 7500 Hz and the high-pass cutoff frequency was set to 1 Hz. In some experiments, a glass microelectrode was used as a comparison. The glass microelectrode was connected to the intracellular A-M Systems amplifier through the compatible headstage and its output was recorded using Axograph X as described in the previous section. Recording obtained using the Intan software had a sampling frequency of 30 kHz. The A-M Systems amplifier used AxoGraph X (AxoGraph Scientific, Foster City, CA) with a sampling frequency of 10 kHz. The time in the two types of recording files were aligned through a common artifact in the recording that occurred whenever stimulating current was injected.

The flex array CFEs and HDCF arrays require different headstages. The flex array was connected to the RHS 32-channel stimulation/recording headstage (M4032; Intan Technologies, Los Angeles, CA). The HDCF array was connected to the RHS 16-channel stimulation/recording headstage (M4016; Intan Technologies, Los Angeles, CA). The extracellular hook electrodes were connected to a modified 18-pin Wire Adapter (B7600; Intan Technologies, Los Angeles, CA), which was attached to a RHS 16-channel stimulation/recording headstage. The headstages for the CFE array (flex or HDCF) and the glass microelectrode were held by hydraulic micromanipulators (MO-203; Narishige, Tokyo, Japan) for fine control of movement.

After desheathing the buccal ganglion and setting up all three headstages (Figure 1E, F), the CFE array was carefully positioned at the surface of the buccal ganglion, oriented to ensure that fibers would penetrate as many neurons as possible (see inset of Figure 1E). The array was then slowly inserted into the neurons of the buccal ganglion using the hydraulic micromanipulators. By looking through the microscope, it was possible to visualize when the tips of the CFEs were within the neuron somata. At the same time, it was generally possible to obtain multiple recordings in different array channels. Nerve BN2-a was then stimulated through the Intan stimulation/recording controller software via the hook electrode to activate motor patterns (2 Hz, 1 ms pulse duration, 300 µA). These motor patterns help to identify the neurons recorded by the array [27]. After the CFE array was positioned, a glass microelectrode was carefully inserted into each neuron that showed recordings in the CFE to confirm that the recordings were intracellular.

To activate a neuron, stimulation parameters were configured in the Intan software. The first protocol was a 2 second biphasic current pulse, with currents ranging from 10 nA to 400 nA. This protocol used a 100% charge balanced current to prevent damage to electrodes when a higher current was required to activate the neuron [29-30].

The second protocol was a 1 second monophasic depolarizing or hyperpolarizing current ranging from 10 nA to 100 nA. This protocol was used to perform comparable stimulation to the monophasic stimulation of the A-M Systems amplifier when a relatively low amount of current was used. To keep the stimulation duration consistent between a CFE stimulation and a glass microelectrode stimulation, an Arduino-based (www.arduino.cc) pulse generator (https://github.com/CWRUChielLab/ArduinoPulseGenerator) was used to drive 1 second long currents in the A-M Systems amplifier.

For both stimulation protocols, the suprathreshold current for stimulation was first determined. To test the effectiveness of the CFE stimulation, the amount of current was stepped by 10 or 50 nA increments to test the response of the neurons to increasing currents. To test the stability of the stimulation, the same amount of current was injected into the neuron multiple times. Between each stimulation, neurons were allowed to recover for at least five minutes.

Impedance values were measured for both the PEDOT:pTS and PtIr coated electrodes. Using the Intan system, alternating current at frequencies of 1000 Hz were passed through the electrodes to determine their impedance. Measurements were done in the following sequence: first, in *Aplsyia* saline; second, after penetrating a neural cell body; third, after injecting current into the neuron (multiple measurements were made each time the neuron was stimulated); and fourth, after the electrode was removed from the neuron and cleaned (see below) and was once again placed in *Aplysia* saline.

To clean the CFEs, the fibers were immersed in a 3% hydrogen peroxide solution for 1 minute to clear any adherent tissue, and then immersed in deionized water for 1 minute. The CFEs were then ready for the next experiment.

### 2.6 Data analysis

Recordings from both systems were plotted after filtering by a second order 3 Hz high-pass Butterworth filter using MATLAB (MathWorks, Natick, MA). This filter was used to eliminate drift and low-frequency fluctuation in the baseline which then made it possible to calculate the action potential amplitude. By comparing the original data and the filtered data, the filtering was found to have minimal influence on the overall shape of the action potentials.

To calculate signal-to-noise ratio, the basic noise level of the recording V_noise_ was determined using an at least 2 s block of non-spiking neural activity. The peak voltage of an action potential waveform was automatically detected by MATLAB using its amplitude. The signal-to-noise ratio (SNR) was calculated by dividing the peak voltage of the waveform of an action potential by the standard deviation of V_noise_ using the equation:

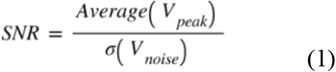

In the multiple channel recordings, the yield of the array was determined by the count of channels in which spiking activity was recorded divided by the total number of array fibers. Because strong firing could result in extracellular recordings on multiple fibers, the counted channels were carefully compared to the actual insertion position of the fibers within the ganglion. Only the fibers that were both in the neuron and showed corresponding spike acitvity were counted as a recording channel.

## 3. Results

### 3.1 CFE intracellular recordings are similar to those from an intracellular glass microelectrode

Previous work has demonstrated that CFEs can be used for chronic extracellular recording in rat motor cortex with a high SNR for up to three months [16] and can potentially record intracellular-like signals [12]. However, intracellular use of CFE arrays has not been tested in detail. To determine the quality of CFE recordings, we compared them to those obtained using a conventional glass microelectrode. Both the CFE and the glass microelectrode were used to impale the same neuron simultaneously, and each electrode’s signal was amplified using a single-channel DC-coupled amplifier.

When a PEDOT:pTS-coated CFE was first inserted into a neuron, a drop in voltage was observed in the recording trace. However, over the course of recording, drift in the baseline of the CFE voltage was larger than in the glass microelectrode recording. Part of the drift may be due to the customized connector attaching the CFE to the intracellular amplifier’s headstage.

Recordings from the two different intracellular electrodes were very similar (Figure 3 A; note schematic to the right indicating the position of the two intracellular electrodes, and the extracellular electrode on the nerve which records from the neuron’s axon in the nerve). Recordings of action potentials at the soma are simultaneous in both intracellular electrodes (Figure 3A, left panel), and the shapes of the action potentials are also very similar (Figure 3 A, expanded time scale, right panel).

**Figure 3.**
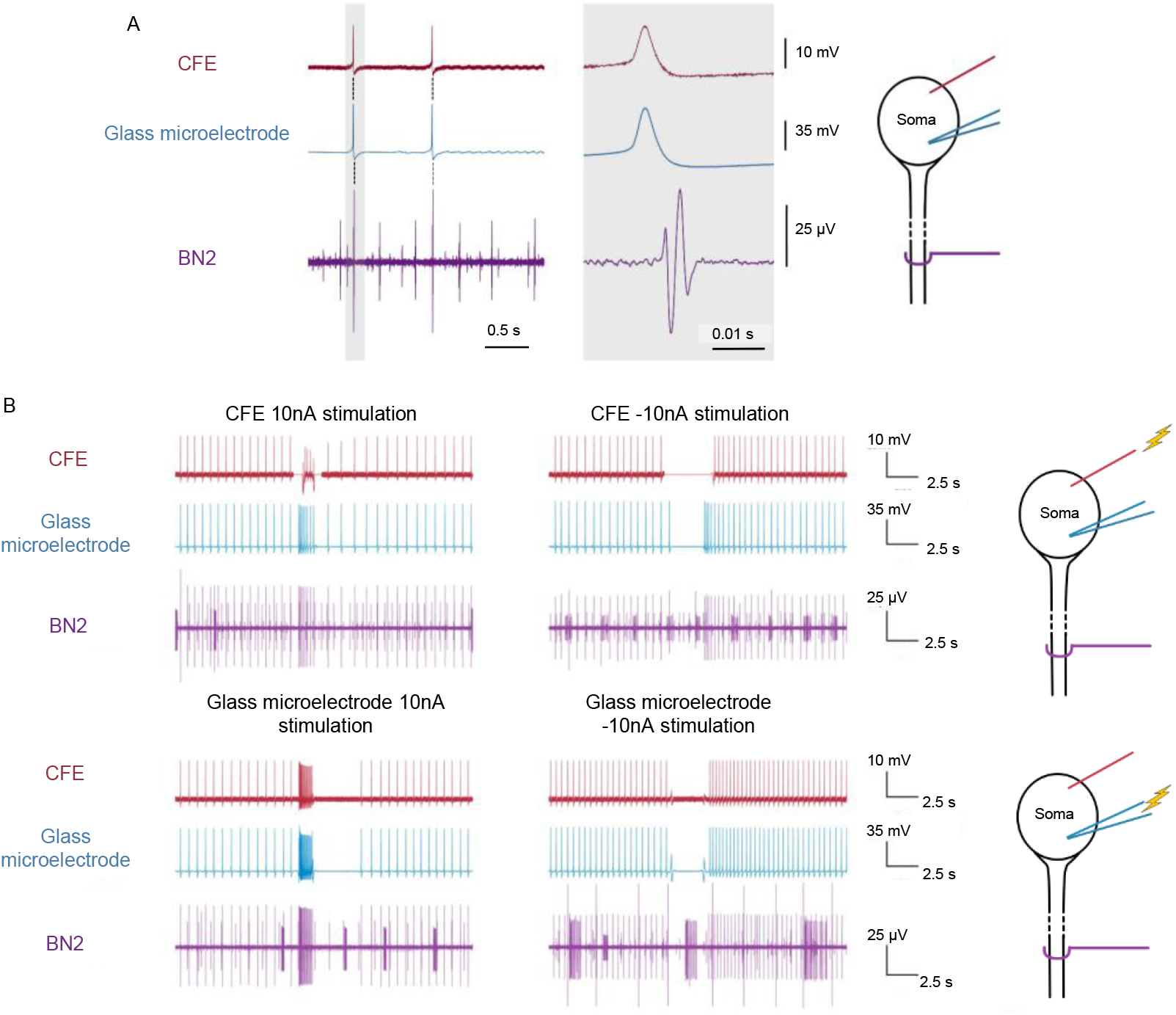
Direct comparison of intracellular recording and stimulation using a sharp glass microelectrode and a CFE, both connected to a DC coupled A-M Systems amplifier. (A) Right panel: Schematic diagram showing insertion of CFE (red line), glass microelectrode (blue), and extracellular recording hook electrode (purple) on the nerve (buccal nerve 2; BN2) containing the neuron’s axons. Left panel: Action potentials recorded at the soma propagated into the axon within BN2. Top trace: CFE recording; Middle trace: Intracellular microelectrode recording; Bottom trace: Extracellular recording of propagating action potential; activity of other neurons that also project on BN2 is visible. Middle panel: A single action potential (marked with a gray bar in the left panel) was expanded in time to show the shape of an individual action potential. In this experiment, the average action potential amplitude recorded by the CFE was 16.47 ± 0.02 mV (mean ± std. dev.) and the average action potential recorded by the glass microelectrode was 59.79 ± 0.03 mV. In this experiment, the SNR of the CFE was calculated as 46, and the SNR of the glass microelectrode was calculated as 318 (see text). (B) CFE current injection compared to a glass microelectrode; both are recorded using the DC-coupled A-M Systems amplifier. Right panel, top: Schematic showing stimulation by CFE, and recording by glass microelectrode and extracellular hook electrode; Right panel, bottom: schematic showing stimulation by glass microelectrode and recording by CFE and extracellular hook electrode. A monophasic pulse of either a 10 nA current (left panels, top and bottom) or a -10 nA current (middle panels, top and bottom) was injected into the neuron through the CFE or glass microelectrode. Action potentials were induced by the injected excitatory current, and spontaneous action potentials in the neuron were inhibited by the injected inhibitory current. During the CFE stimulation, stimulation artifacts were generated and thus replaced by flat lines in the figure. The period of inhibitory current injection was also replaced with a flat line to eliminate the large artifact. Since the action potentials were recorded by the glass microelectrode and projected to BN2, the inhibitory effect of the injected current could still be observed. A postinhibitory rebound was also observed after inhibitory current was injected through the CFE.

The amplitude of the CFE action potential is smaller, and the SNR for the CFE is also reduced. The difference in the signal amplitude may vary with the depth of the CFE electrode insertion and the health status of the neuron after desheathing and electrode insertion. For glass microelectrodes, the recorded amplitudes varied from 7 mV to 60 mV (n = 4 experiments), whereas for CFEs , the recorded amplitude varied from 1.34 mV to 16.47 mV (n = 4 experiments). For each pair of results, the spike amplitude in the CFE was smaller than that in the glass microelectrode. SNR was also calculated to evaluate the recording ability of the CFE. The standard deviation of the noise in the CFE ranged from 0.10 mV to 0.35 mV, whereas the standard deviation of the noise in the glass microelectrodes ranged from 0.17 mV to 0.29 mV, and thus (using equation 1), the SNR for the CFE ranged from 12 to 46, whereas the SNR for the glass microelectrodes ranged from 49 to 318 (n = 4 experiments). These are consistent with some of the recording surface of the CFE remaining outside of the cell, reducing the spike amplitude. The results suggest that PEDOT:pTS-coated CFEs are not as effective as glass microelectrodes for obtaining high SNR recordings, but they can accurately record the shape of intracellular signals as well as a glass microelectrode and have a sufficient SNR to easily distinguish an action potential from the baseline noise.

### 3.2 Intracellular CFE stimulation can activate or inhibit neurons

To understand the dynamics of a neural circuit, the activity of individual neurons should be manipulated, and the effects of this manipulation should be recorded in other neurons. We therefore tested whether neurons could be controlled using carbon fiber electrodes. With a glass microelectrode and a CFE inserted into the soma of the same neuron, identical currents were injected alternately into each electrode to evoke action potentials, and the responses of the neuron to these current injections were compared.

When a PEDOT:pTS-coated CFE was used for stimulation, the injected current successfully elicited action potentials in the neuron which propagated into the neuron’s axon (Figure 3 B, top left panel). During the stimulation, the shape of the action potential was still identical to that recorded by the glass microelectrode. Similarly, inhibitory current injected through the CFE could block spontaneous action potentials (Figure 3 B; top middle panels). At the time that either depolarizing or hyperpolaring current was injected into the CFE, a sudden increase or decrease in the voltage was observed in the CFE recording channel, respectively, before the baseline returned to a normal range. This effect could not be eliminated by adjusting the capacitance using the A-M Systems amplifier. In some cases, the large initial voltage offset could lead to signal saturation, making it difficult to directly record neural activity, but the axonal projection and the glass microelectrode recording still demonstrated the response of the neuron to stimulation.

Similar to a glass microeletcrode, PEDOT:pTS-coated CFEs could activate or inhibit a neuron (Figure 3 B). However, CFEs with the PEDOT:pTS coating do not work well with higher currents (over 60 nA) and are less stable in their impedances after multiple stimulations. Therefore, we switched to the PtIr-coated CFEs for stimulation.

### 3.3 The CFE array can simultaneously record multiple neurons intracellularly

The intracellular A-M Systems amplifier only permits recording from a single channel. To record from a population of neurons and therefore make full use of the arrays, we switched to the Intan system, which could be used for multiple channel recording and stimulation.

During experiments, the array was carefully positioned above the buccal ganglion to reach the maximum possible number of neurons. Once neurons were impaled in a ganglion that had been carefully desheathed, it was possible to obtain stable recordings for at least 4 to 6 hours.

Using the 2×8 flex arrays, on average 65% (n = 5 different experiments) of the channels showed recordings, with 5 - 8 different neurons being recorded simultaneously (for positioning of array relative to the ganglion, see Figure 1 B; results of recording are shown in Figure 4 A). For the flex arrays, since the view of the bottom row of fibers was blocked, it was difficult to determine the actual number of fibers not in neurons.

**Figure 4.**
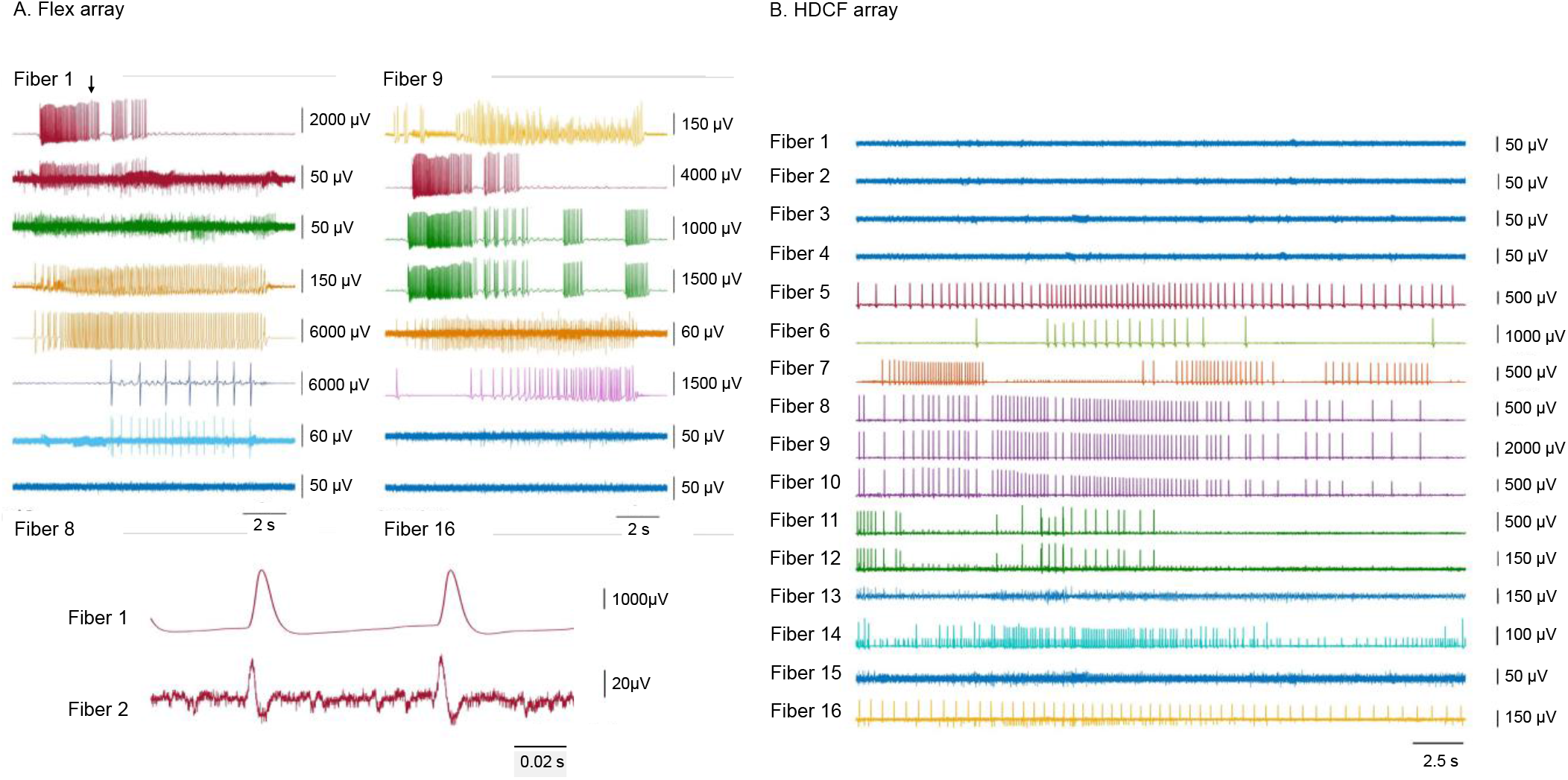
Multiple simultaneous recordings from the flex array (A) or the HDCF array (B) amplifed by the AC-coupled multi-channel Intan system. Recordings that appeared to originate from the same neuron are indicated by the same color. (A) A representative flex array recording showing recordings from seven different neurons in thirteen fibers (1-7 and 9-14) out of sixteen fibers in high potassium saline. The gray bar indicated by an arrow on the top two traces in part A left panels are expanded in time at the bottom to compare the intracellular waveform recorded by fiber 1 and extracellular waveform recorded by fiber 2. (B) A representative HDCF array recording showing recordings from seven different neurons in twelve fibers (5-16) out of sixteen fibers in normal *Aplysia* saline.

Since the HDCF arrays were wider than the ganglion, not all electrodes could be positioned in neurons (Figure 1 D; note that the rightmost electrodes could not be placed within neurons because they are beyond the right edge of the ganglion). During the experiments, we observed that on average 69% (n = 7 experiments) of the HDCF arrays’ fibers were inserted into or had a near proximity to the ganglion, while the rest were in solution. Of the electrodes that could be positioned within neurons of the ganglion, on average 74% (n = 7 experiments) of the channels showed recordings, with 3 - 7 different neurons being recorded simultaneously (Figure 4 B). Since electrodes in the HDCF arrays were all in a single row, it was much easier to determine whether or not they were in a neuron. More generally, the yield of channel recording is related to the positioning of the CFE array, and the size and the natural curvature of the buccal ganglion.

Recordings from the CFEs could be intracellular, extracellular or quasi-intracellular, depending on how deeply the electrode was inserted into the neuron. During CFE insertion, the neurons could be penetrated at different depths because of the natural curvature of a buccal ganglion, causing the signals to be either dominated by intracellular recording or extracellular recording. Therefore, in both flex array recordings and HDCF array recordings, adjacent fibers showed both intracellular recordings and extracellular recordings of the same neuron depending on the penetration depth (Figure 4; for example, in Figure 4 A, the recording on fiber 1 is intracellular, whereas the recording on fiber 2 is extracellular (note the biphasic character of the action potentials in fiber 2)).

To determine whether a recording was fully intracellular or not, a glass microelectrode was inserted into each neuron that had a CFE inserted into it one after the other to compare the shape of recorded action potentials. Fibers that recorded intracellularly generated waveforms that were very similar to those observed from glass microelectrode recordings, even though the two electrodes used different recording systems (Figure 5 A; note the similarity in the recordings on carbon fibers 6 and 11 (top traces in the right panels) to those from a glass microelectrode (bottom traces in the same panels)).

**Figure 5.**
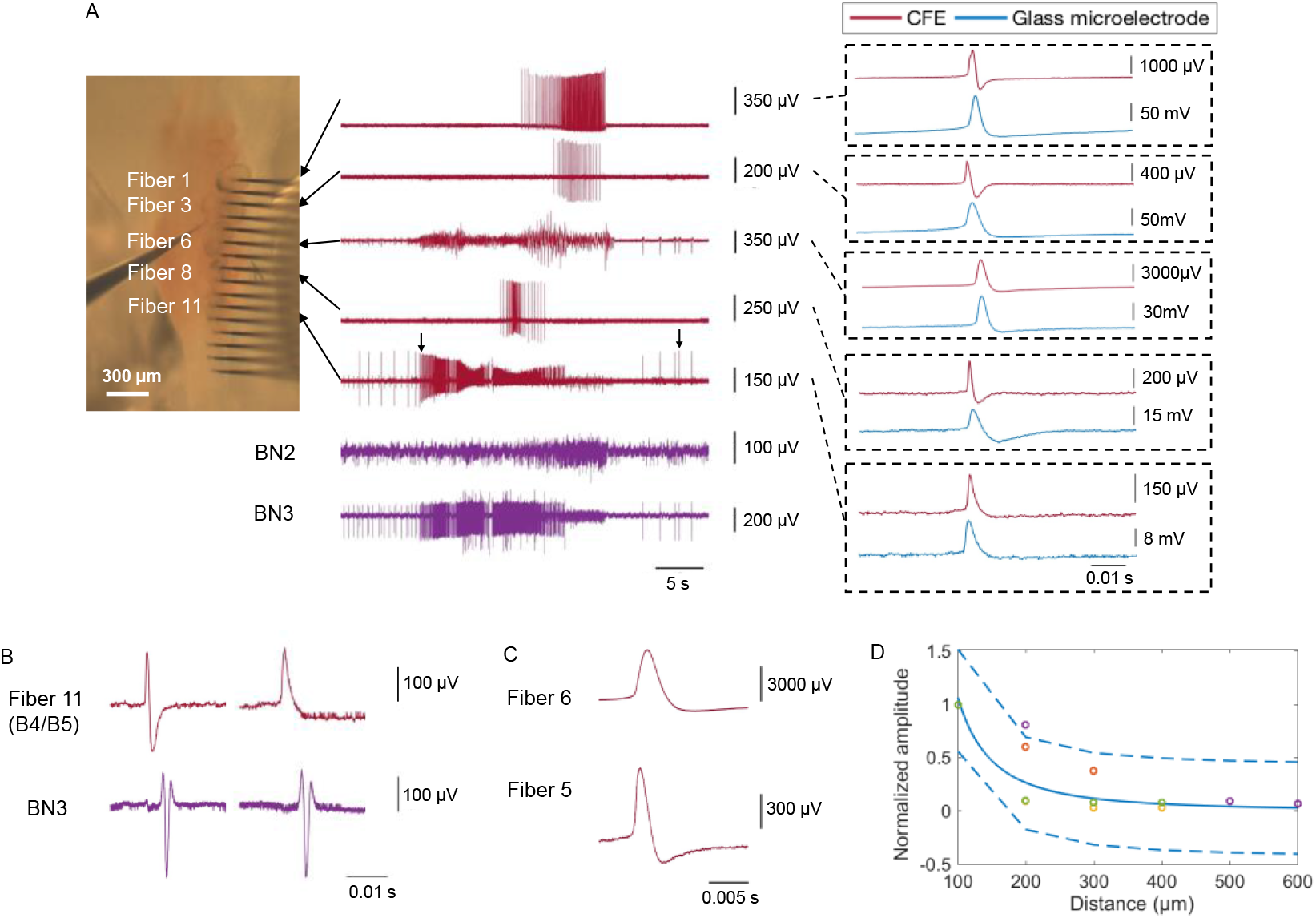
Determination of whether recordings were intracellular or extracellular. (A) A ganglion was penetrated by the HDCF array with PtIr coated tips which was connected to the AC-coupled multi-channel Intan amplifier system. A glass microelectrode connected to the DC-coupled A-M Systems amplifier was then used to penetrate the same neurons in succession to determine whether recordings were intracellular or extracellular. (A) Five different neurons were recorded simultaneously by the HDCF array. From these five recordings, those on fiber 6 and fiber 11 were intracellular recordings, because the shape of their action potentials were very similar to that in the glass microelectrode recording (compare the red traces, recorded by the CFE, to the blue traces, recorded by the intracellular glass microelectrode in each of the boxes on the right). The recordings from the other three fibers were extracellular. Fiber 11 (the fifth trace) recorded from multi-action neurons B4/B5, since the action potentials from these neurons propagated through BN3 as the largest extracellular unit [28]. (B) The insertion of the glass microelectrode into B4/B5 moved the neuron relative to the CFE, so that the recording changed from extracellular to intracellular. Note the change in the shape of the action potential in B4/B5 neuron (top left trace, corresponding to left arrow in part A) from extracellular (biphasic) to intracellular (monophasic, top right trace, corresponding to right arrow in part A). The extracellular recording on the nerve due to the propagating action potential did not change (second trace from top, left and right). (C) Intracellular recording in fiber 6; fiber 5 was adjacent to the cell but was in solution, and recorded an extracellular signal from the same neuron (note change in scale for the two recordings). (D) Percentage fall-off in extracellular spike amplitude as a function of distance. The amplitudes in four trials (shown in different colors) were normalized to the amplitude recorded at the source.The x-axis indicates the distance in µm from the signal source. An inverse square relationship along with a 95% confidence bound was fit to the data points. The latency of these extracellular recordings are on the scale of 0.1 ms. For example, the time latency of the extracellular recording at fiber 9 with a source signal on fiber 11 was 0.47 ± 0.09 ms (mean ± std. dev., n = 67).

Several fibers showed waveforms that were more similar to extracellular recordings when compared to the waveforms on the glass microelectrode (Figure 5 A, recordings from fibers 1, 3 and 8). In addition to lower amplitudes, the depolarizing phase and the repolarizing phase of the extracellular waveform were narrower [31] and the peak occurred earlier than the peak in the intracellular waveform [32]. In one case, inserting the glass microelectrode moved the CFE further into a neuron, changing the recording from extracellular to intracellular (Figure 5 B).

Strong firing in a neuron could also result in extracellular recordings not only in the inserted fibers, but also across the array elements immersed in saline that were adjacent to the ganglion. It was possible to determine the source of the signal by observing the site of the fiber insertion and the highest spike amplitude. Fibers further from the source showed a smaller amplitude (Figure 5 C). As would be predicted from electrical field theory, the amplitude of the extracellular recordings fell off as the inverse square of the distance from the souce (Figure 5 D) [33]. In general, fibers that were recording intracellularly did not pick up extracellular signals from other active neurons, though in some cases, very small extracellular signals could be seen.

### 3.4 Platinum-Iridium coated CFE stimulation was effective and stable

PEDOT:pTS-coated fibers were first used for stimulation. Although it is possible to deliver the current through them (Figure 3 B), the impedance of the PEDOT:pTS-coated fibers was only stable for currents under 30 nA (Figure 6 A, left panel). Higher currents (over 60nA) would quickly increase the impedance and damaged the fibers.

**Figure 6.**
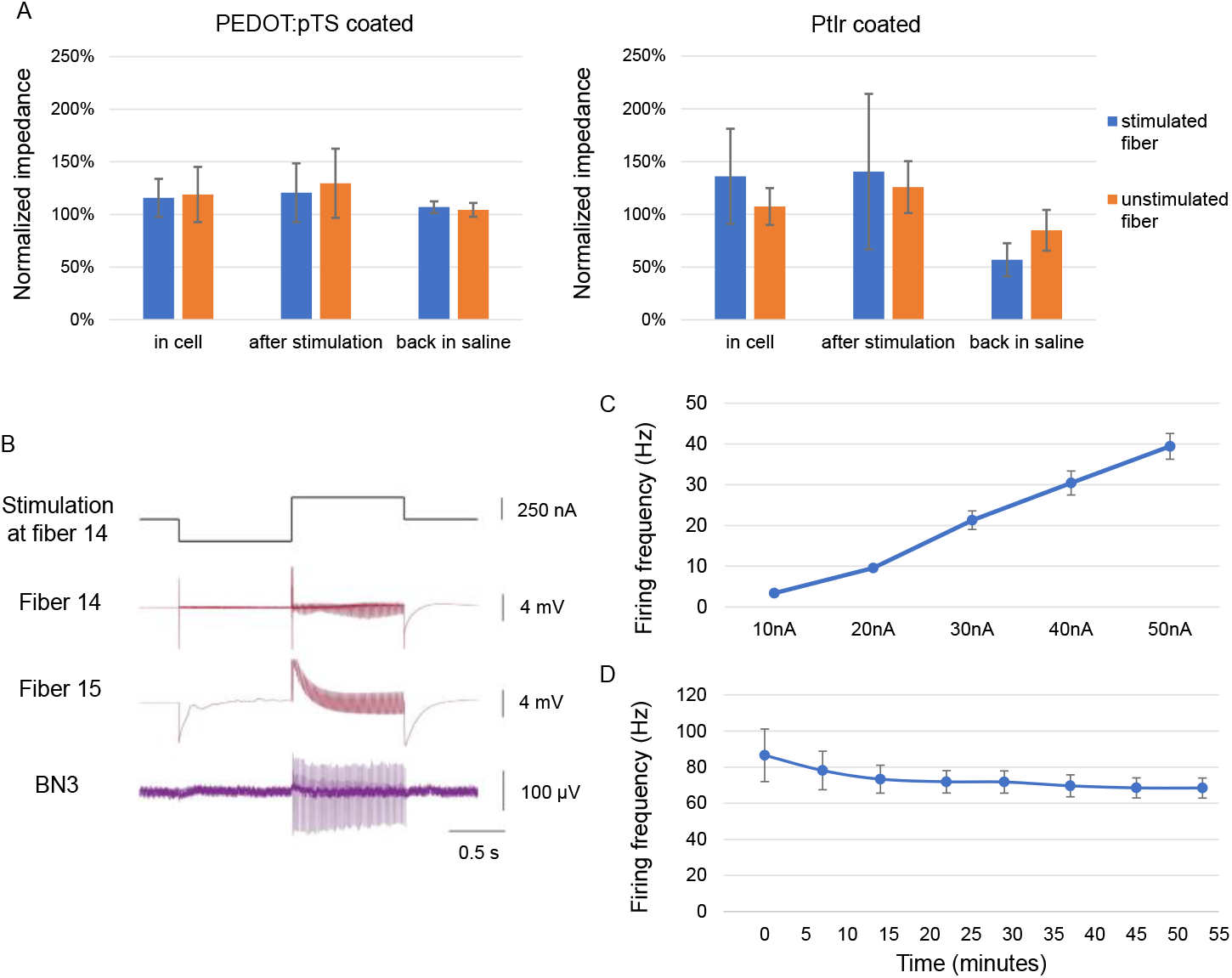
Response to CFE stimulation currents. (A) Normalized impedance to the initial impedance measured in the saline before the cell penetration for PEDOT:pTS-coated and PtIr-coated fibers. After penetrating the soma, electrodes with either coating showed an increase in impedance. PEDOT:pTS-coated fibers had a relatively stable impedance when stimulated with currents up to 30nA. However, the impedance still increased slightly after the electrodes were pulled out of the cell and cleaned to remove any adherent tissue. Higher currents led to very large increases in impedance and damaged the electrodes, and so are not included in these averages. In contrast, PtIr-coated fibers could tolerate currents up to 200 nA without showing signs of damage. After removal from the neuron, these electrodes showed a *decrease* in impedance, even after multiple stimulations with currents of 50-100 nA. Before stimulation (in the cell), the percentage changes in impedance for stimulated and unstimulated fibers were comparable for both PEDOT:pTS- and PtIr-coated fibers (stimulated: 116% ± 18% (n = 6 fibers from 4 experiments) vs. 136% ± 45% (n = 6 fibers from 4 experiments); unstimulated: 119 ± 26% (n = 9 fibers from 4 experiments) vs. 107% ± 17% (n = 9 fibers from 4 experiments) for PEDOT:pTS-vs. PtIr-coated, mean ± std. dev; there are no significant differences between the values for the different coatings in either condition). The middle bars (after stimulation) in the two graphs cannot be compared, because far more net current was passed through the PtIr-coated fibers. Comparing the last set of bars (back in saline) clearly shows that the PtIr-coated fibers could maintain lower impedances over multiple current injections and multiple experiments (stimulated: 107% ± 6% (n = 7 fibers in 5 experiments) vs. 57% ± 16% (n = 6 fibers in 4 experiments); p < 0.0003, t-test; unstimulated: 104% ± 7% (n = 9 fibers in 4 experiments) vs. 85% ± 19% (n = 9 fibers in 4 experiments; p = 0.017; t-test; since 4 t-tests were performed, the criterion for significance should be 0.0125). (B) A neuron activated by a 250 nA current (HDCF array, PtIr-coated tips). Fibers 14 and 15 were inserted into the same neuron, thus recording the same neural activity. 100% charge balanced biphasic currents were injected into the neuron through fiber 14 and recorded by both fibers. The action potentials recorded by the stimulated channel (fiber 14) were somewhat distorted as compared to the action potentials in the recording channel (fiber 15). The action potentials at the soma of this neuron propagated into BN3 with a one-to-one match, indicating that this was the B4/B5 multi-action neuron. (C) Increasing currents generated faster firing rates. 100% charge balanced biphasic currents were injected into the neuron (same pulse waveform as in part B). The stimulation current ranged from 10 nA to 50 nA with a step size of 10 nA. The average firing frequency increased as the current increased. Data are plotted as average ± std. dev., n = {3, 10, 21, 29, 38} spikes. (D) A fixed biphasic current of 250 nA (same pulse waveform as in part B) reliably generated similar output firing frequencies. The average firing rate was plotted against time, and each data point represents a 250 nA current injection after the first trial at time 0. Data are plotted as average ± std. dev., n = {85, 77, 73, 72, 71, 69, 68, 68} spikes.

Since platinum-iridium (PtIr) coating has been shown to have excellent ability to record, low impedance, and a good ability to pass current [14, 30, 34], PtIr-coated CFEs were created and used for intracellular stimulation. They were found to be more effective and stable. Moreover, the impedance of the electrodes *decreased* after stimulations with higher currents than could be applied through the PEDOT:pTS-coated fibers (Figure 6 A, right panel), and, after cleaning, the fibers could be re-used for stimulation in multiple experiments. Thus, our subsequent investigations of intracellular stimulation used the PtIr-coated CFEs.

To investigate stimulation efficacy, different amounts of current were injected into neurons while monitoring the generation of action potentials in axons through extracellular nerve recordings. Action potentials were triggered successfully at the soma (Figure 6 B). The propagation of the action potential through the axon was also observed on the corresponding nerve. Stimulations with higher currents resulted in higher frequency firing (Figure 6 C).

To investigate the stability of stimulation, the same amount of current was repeatedly injected into the neurons, with at least a five minute interval to allow the neuron to recover. Multiple stimulations using the same amount of current consistently activated neurons at a similar firing frequency (Figure 6 D).

The activation currrent threshold of the neurons using PtIr-coated CFE stimulation varied. For glass microelectrodes, the threshold for inducing action potentials in different neurons varied from 10 nA to 100 nA. When using the CFE, in some cases, a current over 200 nA was required to stimulate the neurons; in other cases, currents as low as 10 nA were sufficient. In addition to an innate difference in threshold among different neurons, the different minimum current requirement for neural activation was likely due to fiber penetration depth. When the fiber was intracellular, the stimulation efficiency of a PtIr-coated CFE was similar to that of a glass microelectrode (Figure 7). While both the glass microelectrode and CFE were inserted into the same neuron, the same amount of current was injected into the neuron by either electrode. The axonal projection indicated that action potentials were induced at the soma. The number of action potentials generated by the CFE was slightly lower than but similar to that induced by the glass microelectrode, consistent with the results obtained using the intracellular amplifier for both kinds of electrodes. During the stimulation through a fiber, the shape of the recorded action potential in the stimulated fiber was distorted. In contrast, the shape of the action potential remained unchanged in the glass microelectrode recording as it stimulated the neuron.

**Figure 7.**
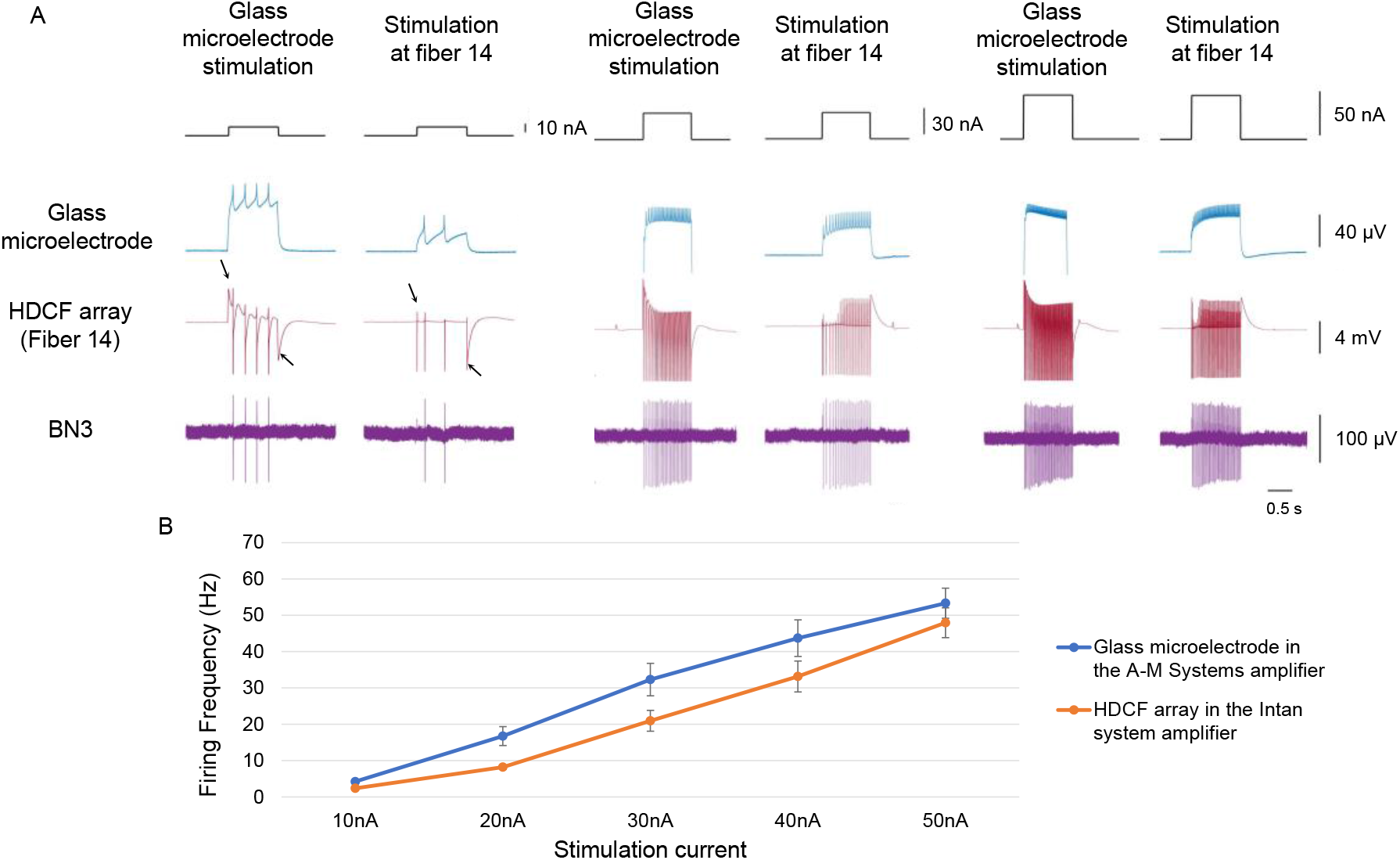
Direct comparison of efficiency of current stimulation using a glass microelectrode and a CFE (HDCF array, PtIr-coated tips). (A) A glass microelectrode and a fiber were inserted into the same neuron. The same amplitude 1-second-long monophasic stimulation currents were sent through either one or the other electrode, and the intracellular response was recorded (arrows indicate the stimulation artifacts; propagating action potentials are observed on the bottom trace, an extracellular nerve recording from BN3). (B) Comparison of the firing rate of the HDCF array’s fiber and the glass microelectrode when the same amount of current was injected (average firing frequency ± std. dev., N = {4, 16, 32, 43, 53} spikes for glass microelectrode and {2, 8, 21, 33, 48} spikes for CFE). Although the HDCF array’s electrode generated lower firing frequencies than did the glass microelectrode, the responses clearly tracked one another.

### 3.5 The CFE could record subthreshold synaptic activity

Subthreshold membrane potentials can regulate neural activity [1-2]. CFEs that record intracellular signals also record subthreshold synaptic activity in the neuron, including inhibitory postsynaptic potentials (IPSPs) and excitatory postsynaptic potentials (EPSPs).

During the experiments, multiple neurons could be recorded simultaneously in the buccal ganglion, including the multi-action neurons that have wide-ranging synaptic outputs to many motor neurons in the buccal ganglion, B4/B5 [19]. With proper positioning of the array, both B4/B5 and some of its synaptic targets in the ganglion could be monitored intracellularly at the same time.

Although the SNR of the CFEs is lower than that of a glass microelectrode, it is sufficiently high to record the IPSPs that were triggered by the action potentials of B4/B5 in its synaptic followers (Figure 8). These subthreshold synaptic potentials were distinguishable from the extracellular recording across the fibers. The subthreshold recordings only occur in the fibers that were recording intracellularly. The time latency of EPSPs (data not shown) and IPSPs was longer than the extracellular recordings on the other fibers. The peak-to-peak time latency was determined to be about 10 ms, which is far slower than the 0.1 ms time latency for extracellular recordings.

**Figure 8.**
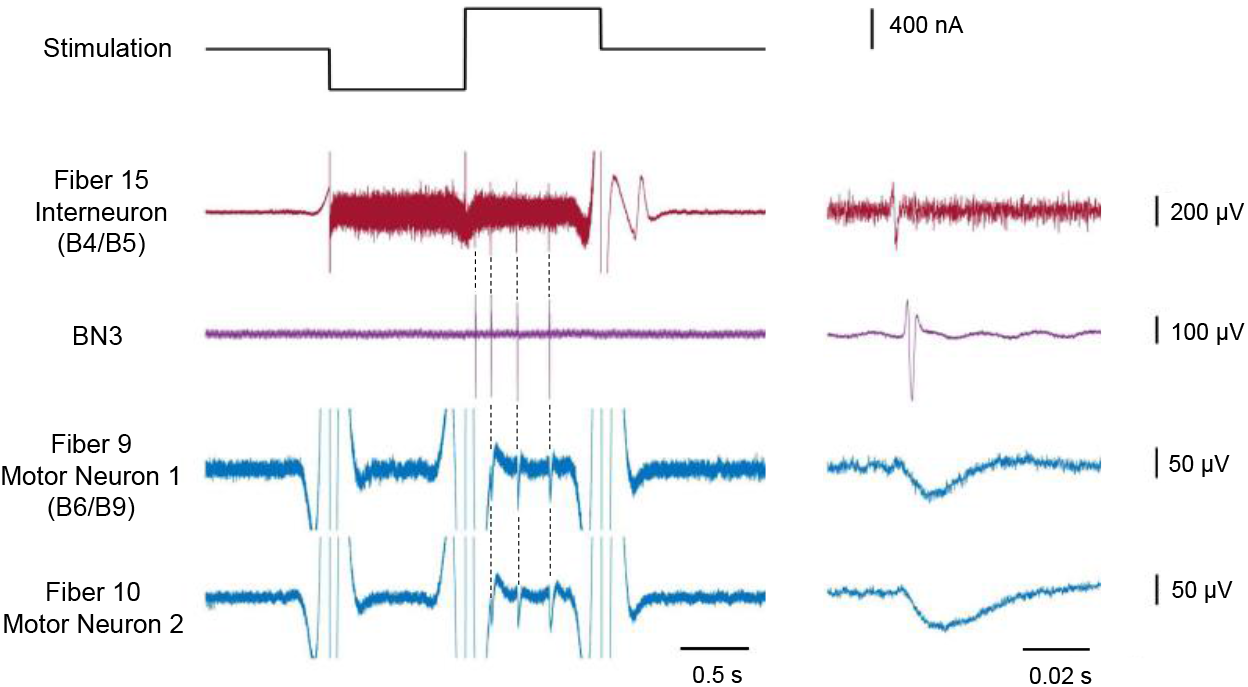
CFEs can record IPSPs induced by stimulation of multi-action neurons B4/B5. The B4/B5 neurons and two motor neurons that are its synaptic followers were recorded simultaneously in a high divalent cation solution that blocks polysynaptic connections. One of the two motor neurons was identified as B6/B9 by its projections on BN2 and BN3 (motor neuron 1). When B4/B5 was activated by a 400 nA biphasic current, IPSPs were observed in the two motor neuron recordings. Right panel shows a temporally expanded version of the record marked by a gray bar in the left panel. Additionally, comparing peak-to-peak time latency, the latency of the IPSPs are on the scale of ∼10 ms. For motor neuron 1, the average latency was 13.6 ± 2.0 ms (mean ± std. dev., n = 9). For motor neuron 2, the average latency was 10.9 ± 1.0 ms (mean ± std. dev., n = 9). This suggests these subthreshold events were not extracellular recordings of B4/B5, which would exhibit a much shorter latency (∼ 0.1 ms). Compare Figure 5.

## 4. Discussion

### 4.1 Summary of Results

Like glass microelectrodes, CFEs can record action potentials (Figure 3 A) and synaptic potentials (Figure 8) with high accuracy. In addition, CFEs can stimulate or inhibit individual neurons (Figures 3 B, 6, 7 and 8) using currents that are comparable to those of glass microelectrodes (Figures 6 and 7). Even though CFEs do not have as low SNR as do glass microelectrodes, and may require more current for stimulation depending on their penetration into a neuron (Figure 7), they can generate stable responses over long periods of time in response to the same current (Figure 6 D). In addition, CFEs make it possible to perform multiple intracellular and extracellular recordings at the same time (Figures 4 and 5), which is much more difficult to do using glass microelectrodes.

### 4.2 Stimulation by PtIr-coated CFEs

Stimulus pulses used during intracellular experiments are of longer duration and lower amplitude, compared to those used for extracellular stimulation. A typical stimulus pulse applied via an extracellular electrode is 0.1 ms - 1 ms and in the range of 10 µA to 10 mA, depending on the application. In contrast, our experiments used pulses that are up to 1 second long and current amplitudes below 1 µA. This difference requires consideration of the charge injection capabilities of the electrodes. Using a 1 second, 100 nA pulse as an example, 0.1 microCoulumbs of charge would be applied via a PtIr-coated CFE. The electrode shown in Fig 2F is approximately 2000 sq microns; thus the charge density of our example pulse is 5 mC/cm^2^. This density is near safe stimulation levels for typical extracellular pulses (short duration, high amplitude). Intracellular pulses (long duration, low amplitude) may allow more complete use of charge storage capacity of a high surface material like PtIr [42]. Nevertheless, high charge density may lead to unacceptable polarization of these electrodes, which may induce local changes in pH or hydrolysis with susequent gaseous O_2_ or H_2_ production [43-44]. However, such negative effects accumulate over continuous pulsing, and a single intracellular pulse may produce harmful reactants at small quantities easily buffered by solution. Future experiments will charactize the electrochemical aspects of intracellular pulses more completely using PtIr electrodes.

### 4.3 Advantages and Limitations of CFEs

Since glass microelectrodes are very stiff, once they are inserted into a neuron’s cell body, it is crucial to keep the entire preparation from moving, or the tip of the glass microelectrode may badly damage the cell membrane. In contrast, CFEs are quite flexible along their lengths, and appeared to tolerate small lateral movements. We were able to obtain stable recordings from CFEs over many hours. Indeed, we anecdotally observed that one effect of movement was to allow the carbon fibers to penetrate the neurons more deeply over time, which improved the quality of the recordings and reduced the current needed to excite or inhibit neurons. Based on these observations, we are confident that CFE arrays may be very useful in semi-intact preparations [20] in which the ganglion may be subject to movement due to attached musculature. Furthermore, the flexibility in response to movement is likely to improve the quality of recordings in chronic implanted CFE-based devices.

The number of neurons that can be recorded using CFEs is already larger than the number that could be easily recorded using standard glass microelectrodes. Even larger numbers of neurons could be recorded by placing several CFE arrays into a ganglion simultaneously. One disadvantage of the current array geometry is that ganglia have curving surfaces, and so arrays need to be positioned at different depths to reach neurons. In future work, it may be possible to incorporate CFEs into soft substrates that can curve and conform to a ganglion’s surface [23], thus increasing the number of neurons that can be recorded simultaneously.

The ability to compare CFE intracellular recordings and glass microelectrode recordings was enhanced by the large soma size of *Aplysia* neurons. Since neuron somata may be much smaller in other animals and in humans, we also performed preliminary experiments on sensory neurons in *Aplysia*, whose diameters range from 20 to 30 µm, which are more similar to those observed in vertebrates and humans. We found that it was also possible to penetrate and obtain stable recordings from these smaller neurons (data not shown). These results suggest that CFEs could be used for a wide range of intracellular recordings in many other nervous systems.

CFEs can clearly be used for simultaneous intracellular and extracellular recordings. Depending on the penetration depth of the CFE, the recordings may have both intracellular and extracellular aspects. The CFE tips were treated by blowtorching to penetrate the cell membrane more easily, but the treatments could create an exposed length of approximately 140 µm at the end [22-23], and this could lead to partial exposure of the conductive part of the CFE to the extracellular fluid. In the future, it may be possible to fabricate tips with much shorter conductive lengths, which could ensure that recordings were completely intracellular. Substituting a sulfuric acid etch for blowtorching would allow the CFE tips to be tailored to a small (< 5 µm in height), sharpened tip [35-36].

It may also be interesting to use the dual ability of CFEs to record extracellularly and intracellularly to intially record from a group of neurons extracellularly, apply spike sorting and circuit analysis techniques that are standard for analyzing such recordings [37-38], and then penetrate the same neurons intracellularly with the same CFEs to provide a “ground truth” for these analysis approaches.

While many novel high channel count neural recording systems are now being deployed [39-41], these are limited to extracellular potentials, where interactions between neurons can only be inferred indirectly. Although the intracellular recordings presented here are shown in invertebrate preparation, the anecdotal report of intracellular-like potentials in a songbird model [12] might suggest that this technique could be optimized for short term intracellular recordings in mammals, using a microdrive. Overall, the ability to achieve multihour intracellular recordings adds a novel and necessary tool for neuroscience studies of circuit interactions.

## Acknowledgements

HJC and YH acknowledge support from NSF Grant IOS1754869; HJC, JPG. PRP, JMR, and CAC acknowledge support from NIH Grant R01 NS118606; PRP and CAC acknowledge support from NIH UF1NS107659; PRP, CAC, JMR, JDW, and EDV acknowledge support from NSF 1707316; and HJC and JPG acknowledge support from NSF Grant DBI2015317. Thanks to the University of Michigan’s Lurie Nanofabrication Facility and Michigan Center for Materials Characterization for the use of their equipment. We are also grateful for helpful comments and suggestions from Dr. Victoria Webster-Wood. Author JDW has a financial interest in PtIr coatings.

## Notes

### Competing Interest Statement

Dr.Weiland has a financial interest in Platinum-iridium coatings.

